# Assessing temporal and geographic contacts across the Adriatic Sea through the analysis of genome-wide data from Southern Italy

**DOI:** 10.1101/2022.02.26.482072

**Authors:** Alessandro Raveane, Ludovica Molinaro, Serena Aneli, Marco Rosario Capodiferro, Linda Ongaro, Nicola Rambaldi Migliore, Sara Soffiati, Teodoro Scarano, Antonio Torroni, Alessandro Achilli, Mario Ventura, Luca Pagani, Cristian Capelli, Anna Olivieri, Francesco Bertolini, Ornella Semino, Francesco Montinaro

## Abstract

Southern Italy was characterised by a complex prehistory that started with different Palaeolithic cultures, later followed by the Neolithic transition and the demic dispersal from the Pontic-Caspian Steppe during the Bronze Age. Archaeological and historical evidence points to demic and cultural influences between Southern Italians and the Balkans, starting with the initial Palaeolithic occupation until historical and modern times. To shed light on the dynamics of these contacts, we analysed a genome-wide SNP dataset of more than 700 individuals from the South Mediterranean area (102 from Southern Italy), combined with ancient DNA from neighbouring areas. Our findings revealed high affinities of South-Eastern Italians with modern Eastern Peloponnesians, and a closer affinity of ancient Greek genomes with those from specific regions of South Italy than modern Greek genomes. The higher similarity could be associated with the presence of a Bronze Age component ultimately originating from the Caucasus and characterised by high frequencies of Iranian and Anatolian Neolithic ancestries. Furthermore, to reveal possible signals of natural selection, we looked for extremely differentiated allele frequencies among Northern and Southern Italy, uncovering putatively adapted SNPs in genes involved in alcohol metabolism, nevi features and immunological traits, such as ALDH2, *NID1* and *CBLB*.

## Introduction

Southern Italy was one of the first European regions to be inhabited by our species. One of the oldest archaeological remains attributed to *Homo sapiens*, has been in fact excavated in Apulia (Grotta del Cavallo ∼45 thousand years ago - kya) and recently associated with the Uluzzian culture (1), also reported in the Middle and Upper Palaeolithic in Greece (2–4), although other interpretations have been proposed (5).

After the Last Glacial Maximum, approximately 20 kya, populations from South-Eastern Europe and West Asia partially replaced continental European Hunter Gatherers (HG) (6,7). Recently, ancient DNA (aDNA) retrieved from remains in North-Eastern Italy post-dated this transition of ∼3 ky, and associated it to the demographic transition occurring between Early to Late Epigravettian in Southern Europe around 17 kya (8,9), as confirmed by archaeological data (10). These observations highlighted the existence of a connection between Western and Eastern Europe well before Neolithic times, with Italy playing the role either of a bridge or a refuge (8).

Neolithization revolutionised the culture and the demography of continental Europe, with Southern Italy being the first colonised place West of the Adriatic Sea (11–13). Recently, the availability of ancient Southern European genomes helped in disentangling the dynamics of the early stages of the Pontic Steppe populations diffusion that occurred in the Bronze Age period (14,15). Differently from the rest of Europe, Greece and Southern Italy appear to have been less impacted by this demic dispersal, being characterised by an additional Iranian-related ancestry (16–19). However, the lack of Southern Italian ancient genomes from the Neolithic period keeps open essential questions regarding this major cultural and demographic transition in the region. Starting from the mid 3^rd^ millennium Before Current Era (BCE) archaeological evidence allows to outline a network of cultural connections interacting along the Adriatic-Ionian axis, operating between two or more different core areas and radiating across trajectories of link and expansion which likely triggered small human groups movements (20,21). As a matter of fact, from about 4.3 to 4 kya, the well-know Cetina-type cultural elements, also related to the Bell Beaker phenomenon in the North-Western Balkans, played an active role spreading from the Dalmatian core area Southwards across the Adriatic in Northern Apulia and South-Eastern Italy also influencing the Ionian Islands and Western Greece (22).

Later on, during the 2^nd^ millennium BCE a flourishing and continuous cultural relationship was established between Southern Italy and Aegean communities especially from the Recent Bronze Age (3.3 to 3.2 kya) onward (23,24). Although the demographic extent of these contacts is not clear, some valuable insights on mobility could be inferred from ceramic crafts. The most recent analytical evidence, relating to the Aegean-type pottery from the core sites of Punta di Zambrone (Tyrrhenian Calabria) and Roca Vecchia (Southern Adriatic Apulia), allow to highlight a strong connection with the Western Greek regions (Ionian Islands, Acarnania, Achaea and Elis) and, to a lesser extent, with Western Crete (25).

Precisely, following the Bronze Age, in a period between the so-called Greek “Dark Ages” (26) and the Archaic Greece (2.7 to 2.5 kya), Southern Italy became a hotspot for the foundation of Greek colonies. The first colonies were set in Campania and Sicily, possibly by settlers arriving from Eastern Greece (Euboea island) around 780 BCE, soon followed by colonies in Calabria, Basilicata and Apulia (27–30), encompassing an area later to be known as *Magna Grecia*. In the second half of the 7^th^ century BCE, some of these settlements, specifically in South-Eastern Sicily (Siracusa and Megara Iblea) and Apulia (Taranto), were attributed to Eastern Peloponnesian founders (31,32).

The nature of the early settlements, the scale of their demographic impact and genetic legacy are still a matter of debate. Some genetic studies (33,34) have tried characterise the demographic impact of these processes in Southern Italy using present-day Italian populations, but none of them had the intent of finely dissecting these ancient components. Furthermore, a recent aDNA study (35) showed that Iron Age Apulians were not yet superimposable to contemporary Southern Italians, pointing to later processes as keys for the understanding of present-day genetic diversity in Italy.

As a consequence of this complex demographic scenario, Italy harbours the largest degree of genetic population structure identified in Europe so far (19,36), making its population a valuable asset for adaptation studies (36–38).

In this study, we performed a genome-wide analysis, unveiling the contemporary genetic structure of present-day South and Eastern European populations, and highlighting recent and past interactions between Southern Italy and Greece. Furthermore, integrating available ancient genomes, we finely characterised the genetic similarities between modern day Southern Italians and different Eurasian ancient groups. Finally, we searched for putatively selected genomic regions by evaluating differences in allele frequencies between Southern and Northern Italians, identifying SNPs potentially under selection that are linked with immunological and dietary traits.

## Results

### Southern Italian genetic structure

Genetic stratification of Eurasian individuals, inferred by *ADMIXTURE* (K = 9), revealed the existence of three main components virtually shared by all the Europeans included in our dataset (Fig. 1A, Fig. S1). These were structured as follows: one (depicted in salmon in Fig. 1A), modal in Sardinia, was also present at high percentages in some Italian and Greek present-day populations; another (yellow), most common in Germany and Slovakia, occurred at similar proportions across Greece, Kosovo and the Italian Peninsula. An additional component (green) is modal in the Caucasus area and also largely present in Italy and Greece.

**Figure 1.**
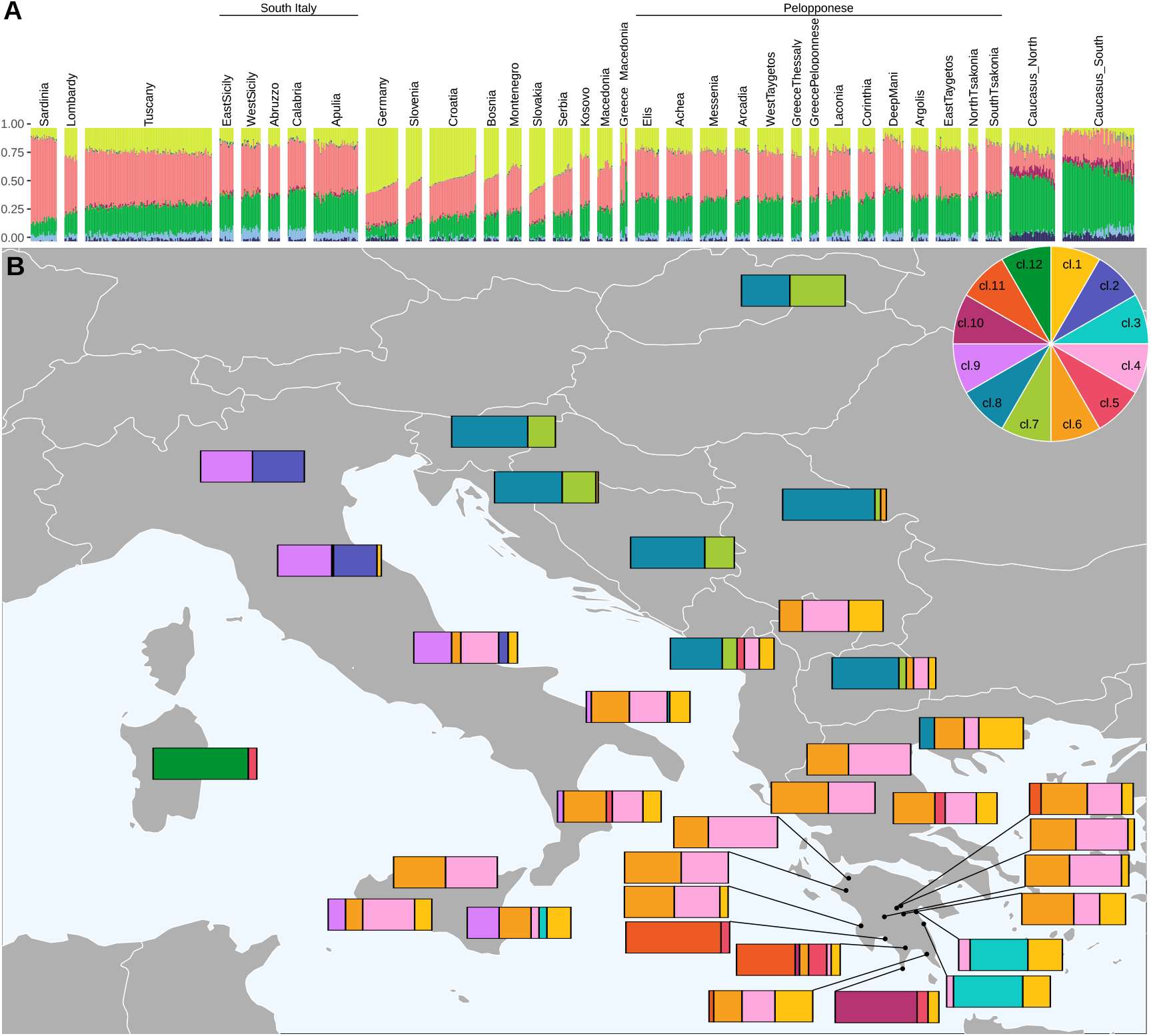
Admixture and genetic relatedness in modern South-Eastern Europeans. A) *ADMIXTURE* plot for K = 9 (lowest CV error) and 34 populations. Populations are ordered according to their geographical longitude. B) Barplots represent the proportion of individuals assigned to a specific cluster (pie in the inset) by the mclust clustering approach (see Fig. S4). Bars are placed on the population geographic origin.

*f4* analysis showed a higher allele sharing of modern Caucasus individuals with present-day Central Europeans than with South-Eastern Europeans, possibly as the result of Bronze Age migrations (Table S2).

The first two PCs in the PCA highlight genetic similarities between Southern Italians and the majority of the analysed Greek populations, with the only exception of the Peloponnesian groups (for sample location see Fig. S2) previously identified as outliers (39), which form a tight cluster also when the third PC is evaluated (Fig. S3).

In order to further dissect the degree of affinity between populations across the Adriatic Sea, we applied a clustering approach on the first ten PCs of a PCA performed on a subset composed by 682 individuals (Fig. 1B, Fig. S4). We detected 12 clusters summarising the genetic similarity among these populations (Fig. S4). Present-day Apulians and Calabrians are characterised by similar profiles, with three clusters (cl. 1, 4, 6) grouping most of their individuals. Western Sicilians mainly fall in cl. 4, which is virtually absent in Eastern Sicily, but it is modal in the Western Peloponnese. Cl. 6 is the second most represented cluster in Apulia, Calabria and Sicily and is found also in Western Peloponnese. Notably, cl. 1 is found at moderate frequencies (from ∼ 16% to ∼ 23%) in the whole Southern Italy, is rare in the Western part of Peloponnese (absent in all Western Peloponnese with the exception of Messenia), but it is present at high proportions on its Eastern part (i.e. Laconia ∼ 36%), Greece-Macedonia (∼ 33%) and Kosovo (∼ 42%). Interestingly, a substantial proportion of Sicilians (between ∼ 16 to ∼ 30%), are part of a cluster which includes Northern and Central Italian areas (cl. 9), and is absent in Greece and Peloponnese. East Sicily and Apulia are the only Italian groups with individuals assigned to a specific cluster observed in drifted Tsakonians (North and South) (cl. 3). Similarly, one individual (out of 17) from Calabria and two Sardinians (out of 24) were assigned to a cluster (cl. 5) composed mainly of the Deep Mani and Taygetos samples, which were previously described as populations that experienced genetic drift (39).

### Modelling the relationship between modern Southern Italian and ancient Eurasians

We evaluated the genetic variation in Southern Italy with respect to ancient groups assembling a dataset composed of 138 ancient Eurasian individuals dating between the Palaeolithic and the Iron Age.

Ancient genetic profiles were preliminarily investigated through PCA by projecting ancient genotypes onto the first two eigenvectors inferred on modern individuals (Fig. S5). Hunter Gatherers individuals from West (WHG) and East (EHG) Europe cluster on the right side of the PCA delineating a West to East cline along the second PC. Most of the individuals enriched in “Neolithic ancestry” are placed close to the genetic variability of present-day Sardinians. This group of samples also includes more recent individuals (from Late Neolithic to Copper Age) who in turn are scattered close to present-day inhabitants of the island (Fig. S5A). Interestingly, the only Greek Neolithic individual is close to Sardinians and European Early Neolithic samples and is part of a tight cluster that includes Anatolians and two Peloponnesians, from the Neolithic Age. On the other hand, three out of the five Neolithic Peloponnesians, together with the totality of Minoans and Mycenaeans included in our dataset plot towards the genetic variability of people currently inhabiting Southern Peloponnese (Maniots and Tsakonians) and Southern Italian regions (Sicily, Calabria and Apulia) (Fig. S5B). Modern Southern Italians are closer to Southern European Neolithic and Bronze Age samples (Neolithic Peloponnesians and Minoans) than most modern Peloponnesian groups, with the exception of the Deep Mani and Taygetos individuals (Fig. S5B). The affinity between Southern Italians and ancient samples was also investigated by the *f-statistics*. First, we tested the affinity symmetry of Northern Italians and other Italian groups (OIGs) with respect to Anatolia Neolithic (AN) samples (OIGs). All the *f4* (Mbuti, AN, OIG, Lombardy) show a significantly higher affinity between Lombardy and AN, with the only exception of Sardinia (Table 1). This pattern is also evident when Bronze Age Greeks (Table 1) or ancient Steppe groups (Table S3) are used in place of AN. These observations suggest that populations from Central and Southern Italy had a lower contribution from AN than Lombardy, or alternatively, that Central and Southern Italians received contributions from other different groups, possibly associated with present-day Middle Eastern or African regions (19,40,41).

**Table 1.** *f4-outgroup* statistics to compare the relationships between Neolithic and Greek Bronze Age samples with Northern Italians and Sardinians.

| Outgroup | Ancient Sources | OIG<sup>1</sup> | North Italy | f4 | SD | Z |
| --- | --- | --- | --- | --- | --- | --- |
| Mbuti | Anatolia_N | Abruzzo | Lombardy | 0.000591 | 0.000196 | 3.02 |
|  |  | Calabria |  | 0.001111 | 0.000172 | 6.442 |
|  |  | EastSicily |  | 0.001381 | 0.000177 | 7.789 |
|  |  | Apulia |  | 0.000915 | 0.000153 | 5.995 |
|  |  | Sardinia |  | -0.001415 | 0.000177 | -7.98 |
|  |  | Sicily |  | 0.001393 | 0.000349 | 3.992 |
|  |  | Tuscany |  | 0.000436 | 0.000138 | 3.172 |
|  |  | WestSicily |  | 0.001421 | 0.000171 | 8.321 |
|  | Minoan | Abruzzo |  | 0.000839 | 0.000235 | 3.573 |
|  |  | Calabria |  | 0.001392 | 0.000212 | 6.576 |
|  |  | EastSicily |  | 0.001676 | 0.000218 | 7.683 |
|  |  | Apulia |  | 0.001016 | 0.000183 | 5.562 |
|  |  | Sardinia |  | -0.000835 | 0.000215 | -3.884 |
|  |  | Sicily |  | 0.001343 | 0.000433 | 3.099 |
|  |  | Tuscany |  | 0.000582 | 0.00017 | 3.417 |
|  |  | WestSicily |  | 0.001574 | 0.000206 | 7.628 |
|  | Mycenaean | Abruzzo |  | 0.00077 | 0.000372 | 2.068 |
|  |  | Calabria |  | 0.000832 | 0.000335 | 2.486 |
|  |  | EastSicily |  | 0.001749 | 0.000348 | 5.025 |
|  |  | Apulia |  | 0.001128 | 0.000299 | 3.772 |
|  |  | Sardinia |  | -0.000666 | 0.000339 | -1.968 |
|  |  | Sicily |  | 0.002023 | 0.000672 | 3.008 |
|  |  | Tuscany |  | 0.000668 | 0.00028 | 2.387 |
|  |  | WestSicily |  | 0.00142 | 0.000344 | 4.13 |

<sup>1</sup>Other Italian Group

However, when the affinity of Italian groups with African and Middle Eastern populations was tested, Southern Italians resulted not significantly closer to any of the two (Table S4).

Except for East Sicily and Calabria, no significant results were detected for the combination *f4* (Mbuti, Steppe, OIG, DeepMani), confirming a similar genetic composition for Maniots and Southern Italian groups. In contrast, replacing Deep Mani with other groups from different Peloponnesian areas resulted in the latter showing a higher affinity to Steppe ancestry than present-day Southern Italians (Table S3).

We further explored the admixture model of Southern Italian groups using *qpAdm*, which fits a vector of *f4* values by summarising the relationships of test and source groups (left populations) to a set of right populations. First, we used the main ancestral populations of present-day Europeans (Anatolia_N, WHG, Iran ChL and EHG) (14,42) as putative sources (Fig. 2A). Notably, all the tested populations, with the exception of Sardinians and Greece Macedonians, show remarkably similar proportions of Anatolia Neolithic source, suggesting that any difference in ancestry observed with the *f4-outgroup* statistics is very subtle. Moreover, most of the tested populations but Sardinians show a relatively high proportion (from ∼ 28 to 54 %) of ancestry related to samples inhabiting Iran in the Chalcolithic period (Fig. 2A). Generally, similar AN ancestral proportions were observed when a model including AN, Steppe_EMBA and Iran ChL was considered (Fig. 2B). In this setting, the Northern Italian group (Lombardy) presents a higher contribution of both AN and Steppe EMBA than the Southern Italians, where a remarkable contribution of Iran Chalcolithic was detected (Fig. 2B).

**Figure 2.**
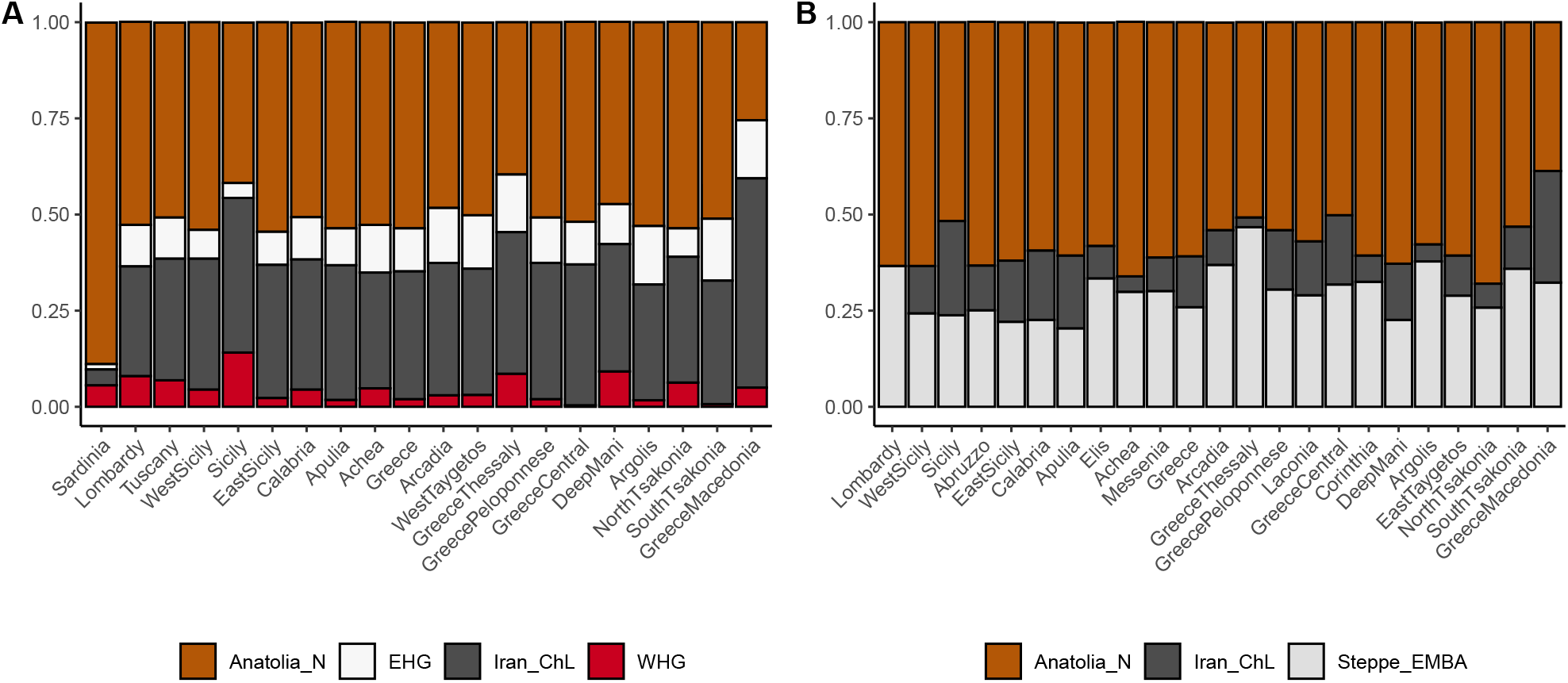
Modelling of ancestry proportion in South-Eastern European populations. A). Supported *qpAdm* proportions for the main ancestral populations of present-day Europeans. B) as in A) using a Steppe EMBA ancestry instead of WHG and EHG ancestral groups as indicated in the legend. Colours are as in Fig. S5.

The admixture graph (Fig. 3) helped in delineating ancestral genetic population differences and similarities between present-day Southern Italians and Greeks in whom an overall higher Steppe-related ancestry was detected (Fig. 2B, Table S3). In particular, we were able to fit a graph admixture model in which 63% of the Minoan genome was found in a population that contributed 97% to the genetic pool of present-day Apulians (Fig. 3A). Contrary, modern Greeks were successfully modelled as an admixture of populations related to present-day Balkan populations (Serbian, (43)) and Bronze Age Greek groups, here represented by Minoans (Fig. 3B). This could potentially explain the differences found in Steppe-related ancestries between the two present-day populations. Additionally, these results suggest a more recent gene exchange between Greeks and neighbouring Slavic-related populations. Alternatively, they could be explained by different ancient legacies between the two sides of the Adriatic Sea, as the result of pre-Bronze Age genetic structure in the Balkans.

**Figure 3.**
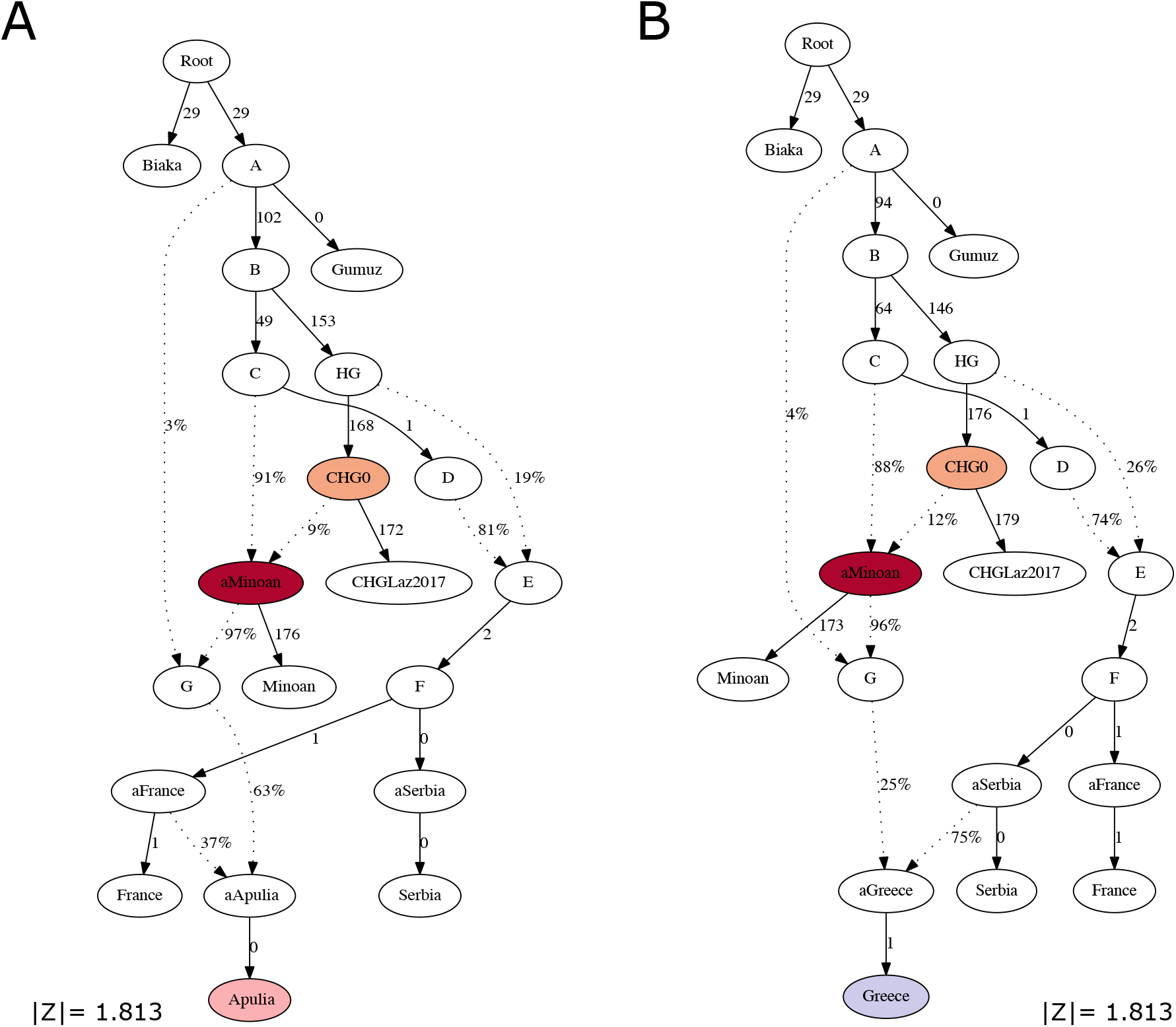
Graph modelling of present-day Apulians and Greeks. Z scores of both trees are between −3 and 3. Ancestries highlighted in shades of red are ancestries specifically shared with Bronze Age Greeks. Letters from A to F refer to unsampled ancestral populations.

### Investigating putative signature of adaptation among Northern and Southern Italians

Following the differences in ancient ancestry contributions across the Italian Peninsula, we applied the Population Branch Statistics (PBS) analysis in order to identify markers showing significant allele frequency differences among pairs of Italian populations using CHB as an outgroup. We estimated the average PBS across windows of five consecutive SNPs and focused on the top ten windows with higher frequency in Northern (positive PBS values) and Southern Italy (negative PBS values), respectively (Fig. S6). We finally annotated the variants with the most extreme PBS values (above the 99.5 percentile and below 0.05) and evaluated if they were enriched for any biological or functional feature.

Among the top 20 windows, seven do not overlap with any known gene (Fig. 4, Table S5). Of those showing the highest allele frequencies in Northern Italy, one encompasses *PABPC4L* (*H. sapiens* poly(A) binding protein, cytoplasmic 4-like), a gene associated with venous thromboembolism (44), birth weight (45), blood protein level and alcohol consumption (46); another, located on chromosome 1, encompasses the gene *NID1* (Nidogen 1), which is associated with blood proteins (47) and platelet levels (48,49), and a third on chromosome 3 maps within the *CBLB* (Cbl proto-oncogene, E3 ubiquitin protein ligase B) gene, involved in the immune response by regulating T- and B-cell receptors (50). Of the windows with extreme negative PBS values, the two most divergent are located within chromosome 12 and encompass 15 genes (Fig. 4, Table S5). They include *ALDH2*, known to affect the metabolism of alcohol and correlated to different behavioural traits associated with alcohol consumption (51). We note here that our dataset does not contain SNPs within this gene, and it is thus possible that this signal is led by variants in other regions. Finally, we found extreme divergence in a region within the *EDAR* (Ectodysplasin A receptor) gene, associated with variations in head hair morphology and facial hair thickness (52). Although the SNP rs3827760 (chr2:109513601), affecting hair follicle thickness and shape, is not present in our dataset, it is unlikely that it is driving the adaptation signal, as also observed in a previous exome analysis (37).

**Figure 4.**
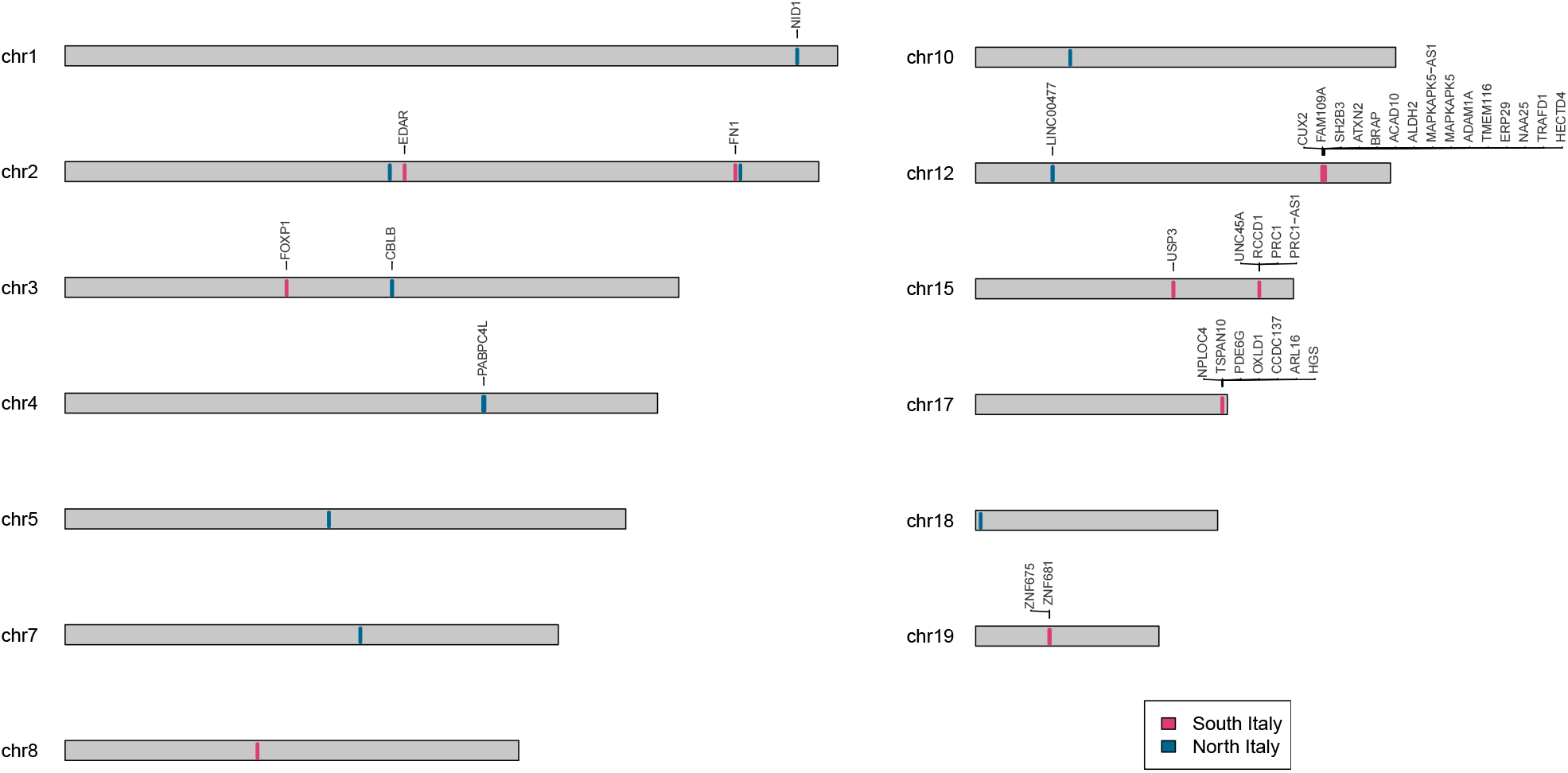
Karyotype plot of the top 10 windows from PBS analysis. In red and blue are indicated the chromosome locations for the top 10 PBS in North and South Italy, respectively. Where present, the set of annotated genes is presented.

## Discussion

Since prehistoric times Southern Italy has been a cross-roads of many human groups and cultures, and their genetic signatures are still present in the DNA of individuals currently living in this area (19,53–55).

In this study, we highlighted a high similarity between Southern Italy and the Peloponnese. In fact, our cluster analysis showed that present-day South-Eastern Peloponnesian populations have high genetic affinity with modern Apulians, Calabrians and South-Eastern Sicilians, all characterised by a cluster composition different from those displayed by other Greek groups (Fig. 1B, Fig. S3). Additionally, individuals from Western Sicily show similarities with populations inhabiting the Western part of Peloponnese (Fig. 1B, Fig. S4). Although establishing the chronological context for this affinity using present-day genomes might be challenging, our results are in accordance with archaeological and historical sources that attributed the origin of Greek colonies in South-Eastern Sicily and Apulia from populations inhabiting the southern and Eastern parts of the Peloponnese (31,32). Uniparental Y-chromosome findings are also in agreement with these observations revealing Eastern Peloponnesian ancestries in East Sicily (34) and shared haplogroups among modern-day Greeks and populations living in Southern Italian areas colonised by Greeks such as the Salento (Apulia) and the Ionian coast of Calabria (56). The lower affinity with other Balkan populations could be attributed to a lower influence by inland populations, such as Slavic-related people (57) and/or genetic drift in Tsakones and Maniots as suggested by historical sources (39). Therefore, our results imply a high affinity between Southern Italians and Peloponnesians possibly abrupted very recently by major events of migrations and/or admixture as the one recorded during the Middle Age period (58). However the observation that, in some analyses, Southern Italians and ancient Greeks share more alleles than modern and ancient Peloponnesians, may suggest a scenario including the preservation of an ancient population signal in the genome of Southern Italians that was likely diluted by inland migrations in Greece (Fig. 3).

Notably, a substantial genetic affinity was highlighted between Peloponnesian individuals from the Neolithic and the Bronze Age, whose signatures are also recorded in Italy. Recent genomic studies (19,33,38) detected a contribution ultimately derived from the Caucasus (CHG) in modern Southern Italians and likely brought in Italy as early as the Bronze Age, although their demographic dynamics are still not known. Overall, these results are in agreement with the detection of a small proportion of Iranian-related ancestry in Sicilian Middle Bronze Age samples (17), which could be tentatively linked to the spread of the Mycenaean culture (59). Interestingly, our results modelled the source of this contribution as a mixture of AN and Iran Chalcolithic ancestries (14). The latter was found consistently across Southern Italy and the Peloponnese, confirming again common genetic sources shared between these two regions (Fig. 2).

The largest degree of genetic heterogeneity across the European continent has been so far recorded in the Italian Peninsula (19), an important aspect to consider when epidemiological and translational studies in Italy are planned. Recent researches taking advantage of high coverage whole-genome and whole-exome sequences linked variants under selection in Italians to genes that were related to insulin secretion, obesity, thermogenesis, alcohol consumption, pathogenic-response, skin colour and cancer (37,38). Here we assessed the occurrence of extreme differences in allele frequencies between Northern and Southern Italy close to the *ALDH2, NID1*, and *CBLB* genes that may have a role in alcohol metabolism, nevi features and immunological traits, respectively. Although the analysed dataset contained only ∼100k SNPs, this replication in the possibly affected phenotypes may suggest an actual important difference among groups on the two parts of the peninsula. Nevertheless, genomic scans are sometimes controversial and difficult to interpret, and a deeper investigation, possibly involving multiple tests on denser datasets, is highly desirable. In conclusion, we provided new insights on the composition of modern genomes of individuals collected from Southern Italy, a still poorly sampled region of Europe. The fine characterisation of the time of arrival and routes of migrations related to the detected ancient signals is expected to be addressed only by the analysis of Apulian ancient genomes covering a period ranging from the Late Neolithic to the Early and Middle Bronze Age periods. The differences identified in modern-day Southern and Northern Italians have significant phenotypic implications and call for more extensive investigations on a larger number of markers and individuals.

## Material and Methods

### Dataset

We assembled a dataset comprehensive of 2,662 present-day individuals from the Mediterranean basin, Europe and Africa genotyped with different Illumina SNP arrays (39,43,57,60–68) (Table S1).

Only SNPs and individuals with a missingness rate lower than 1% were retained (*--geno* and *--mind* flags equal to 0.01). Pairs of related individuals (PI_HAT > 0.05) were identified using PLINK1.9 and within each pair one was randomly removed (69). After merging and pruning, we retained 93,647 linked and 56,956 unlinked (*--indep-pairwise* 100 25 0.2) SNPs. The modern-day dataset was merged with 138 publicly available ancient individuals (6,14,15,18,42,70–82), for whom genotypes of at least 20,000 markers were available (Table S1).

### Genetic structure analyses

#### Admixture analysis

The present-day dataset was processed using *ADMIXTURE* (83). Ten independent runs of unsupervised *ADMIXTURE* for different numbers of clusters (K), from 2 to 20, were performed. Runs were summarised, ordered and aligned using *CLUMPAK* (84). The lowest *Cross Validation* (CV) median error across all the independent runs was observed for K= 9.

#### Principal component analysis (PCA)

Principal Component Analysis was performed to evaluate the genetic structure of Southern Italians in the context of other present-day and ancient Eurasian individuals. All the PCAs were performed with the *smartpca EIGENSOFT* package (85).

First, we considered a dataset including 826 present-day Europeans. Three iterations were initially used to remove outliers resulting in a final dataset of 804 individuals. Secondly, we projected 138 ancient samples on the PCs estimated from the genetic variability of 1,165 present-day Western Eurasians using the *lsqproject* and *autoshrink* options set to *YES* and five iterations to remove outliers.

#### Genetic clustering of Mediterranean populations

In order to group homogenous individual at a more detailed resolution that that offered by *ADMIXTURE* or by the first two PCAs, we applied the clustering approach to finite mixture models implemented in the *mclust* R package (86) using 36 as the number of mixture components for which the BIC is calculated. The approach was performed on the first 10 principal components estimated on 682 present-day individuals from Eastern regions of the Mediterranean basin using *smartpca* and five iterations to remove outliers. The highest BIC value was reported for 12 clusters.

### F-statistics

*f3* and *f4* outgroup analyses were performed using *admixr* v0.7.1 R package (87) which exploits *ADMIXTOOLS v4*.*2* package (88).

We preliminary explored both Denisovan (89) and the Mbuti, a rainforest African group of hunter-gatherers, as outgroups; since no differences were encountered, we selected Mbuti as the outgroup. A |Z| score higher than 3 was considered significant.

### Ancient admixture analyses

*QpGraph* models were generated using *ADMIXTOOLS v6*.*5* (88) with Mbuti or Japanese set as outgroups. We used the default settings and filtered out results with a maximum fit for f-statistics (|Z| score) lower than 3.

*QpWave* and *qpAdm* in *ADMIXTOOLSv4*.*1* package were used as in (19,90) by testing all the possible combinations of two, three and four sources on a set of target populations. We firstly used *qpWave* to establish, using *f-statistics* and a set of outgroups, whether the tested sources were significantly different and could be used to reconstruct the ancestry of a target population. Subsequently, *qpAdm* was employed to infer the proportions of the tested sources.

The threshold used for *qpWave* to establish the significance was set at 0.01, while the set of sources included: Minoan, Mycenaean, Iran Neolithic (Iran_N), Anatolia Bronze Age (Anatolia_BA), Anatolia Neolithic (Anatolia_N), Steppe Early Middle Bronze Age (Steppe_EMBA), Western Hunter Gatherer (WHG), Eastern Hunter Gatherer (EHG), Caucasus Hunter Gatherer (CHG), Peloponnese_Neolithic, Late Western Caucasus Bronze Age culture (LateMaykop), Iran_ChL (Chalcholitic), SteppeMaykopoutlier, Morocco (Table S1).

The set of outgroups comprised: Mota, AfontovaGora3, EHG, CHG, WHG, Goyet, Vestonice16, Malta1, ElMiron, Iran Neolithic (Iran_N), Levant Neolithic (Levant_N), Anatolia Neolithic (Anatolia_N), Kostenki14, Natufian, Ust_Ishim (Table S1).

### Selection scan

The Population Branching Statistic (PBS), which harnesses the differences in pairwise *F*_*st*_ values between three populations, was estimated with the function *allel*.*pbs* of the *scikit-allel python* package (https://zenodo.org/badge/latestdoi/7890/cggh/scikit-allel) using a window_size of 1 and setting the *normed* option as *True*, which normalised the PBS as in (91). We focused on differences between Southern (Abruzzo, Calabria, East Sicily, West Sicily, Sicily, Apulia) and Northern Italy (from Lombardy, Tuscany) using the Han Chinese in Beijing (CHB) population as an outgroup. We averaged the PBS score across windows comprising five consecutive SNPs. Gene annotation was performed with CADD (Combined Annotation Dependent Depletion) (92) and known genetic association for each variant was assessed interrogating the GWAS catalogue, considaering only associations with a *p-value*□<□5E^−08^ (retrieved on November 2021; (93,94)).

