## Supplementary Figures for "Assessing temporal and geographic contacts across the Adriatic Sea through the analysis of genome-wide data from Southern Italy"

### Supplementary Information

#### Supplementary Figures

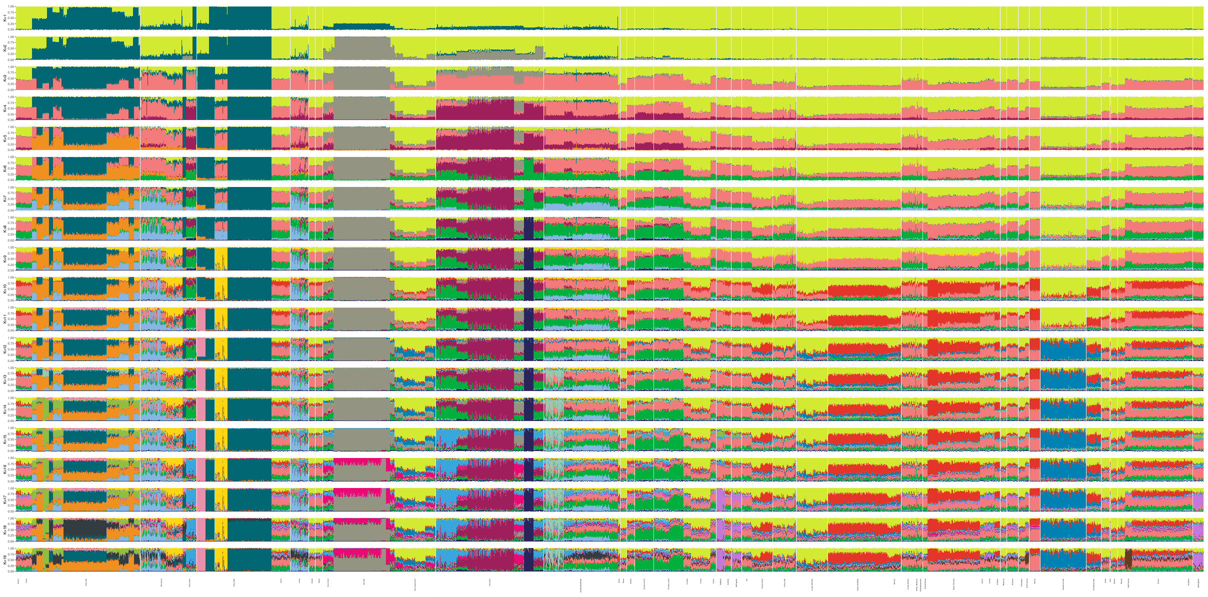

**Fig. S1**. **ADMIXTURE results.** ADMIXTURE plot summarising 10 runs from K=2 to K=20 of all the 2,622 present-day individuals present in the dataset and grouped in 55 macro-areas.

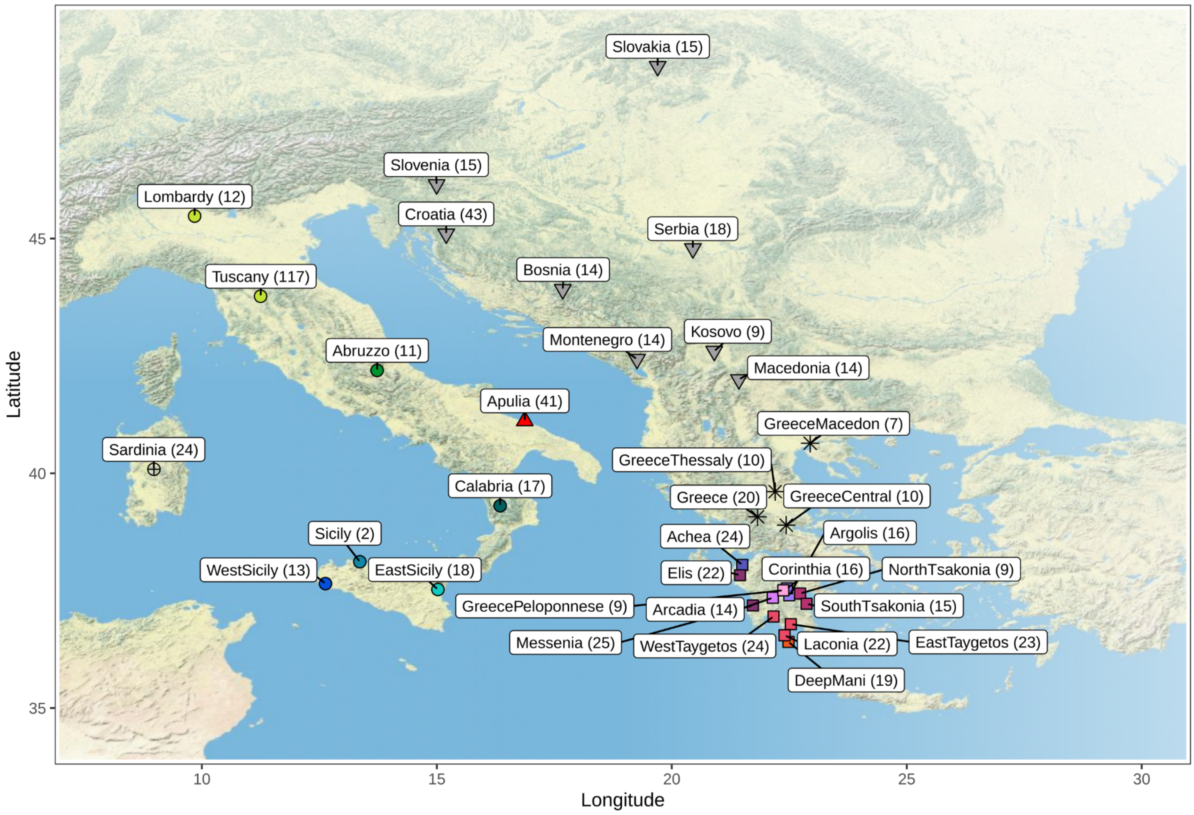


**Fig. S2**. **Sample location of present-day Eastern Mediterranean Basin populations.** Symbols coded as in the PCAs (Fig. S3 and Fig. S5).

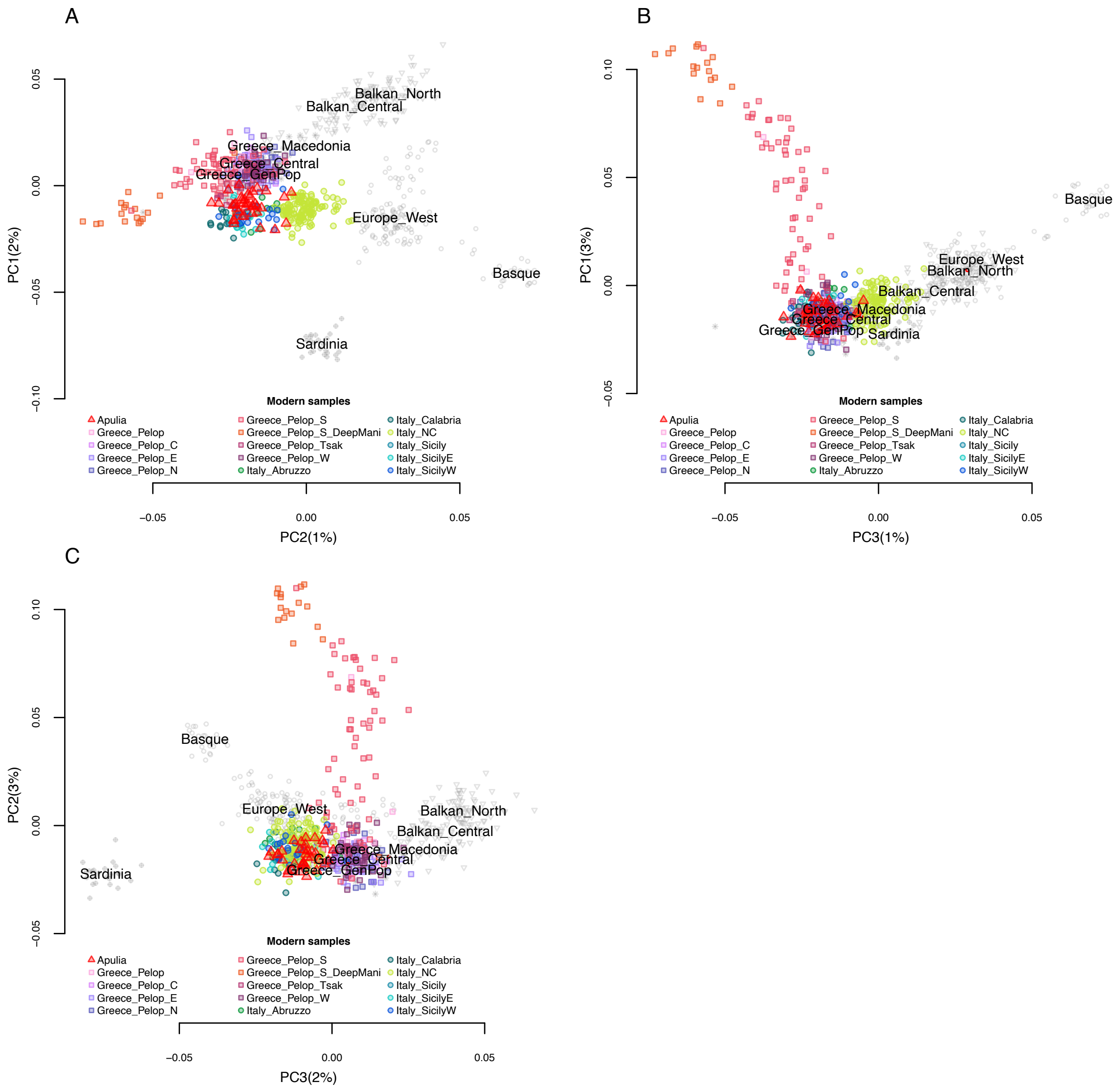


**Fig. S3**. **Genetic structure in present-day Eastern Mediterranean Basin populations in the context of present-day Europeans.** Principal components plot, each dot represents an individual colour-coded according to the populations shown in legend. (A) PC1 on Y axis and PC2 on X axis; (B) PC1 on Y axis and PC3 on X axis; (C) PC2 on Y axis and PC3 on X axis. Label codes can be found in Table S1.

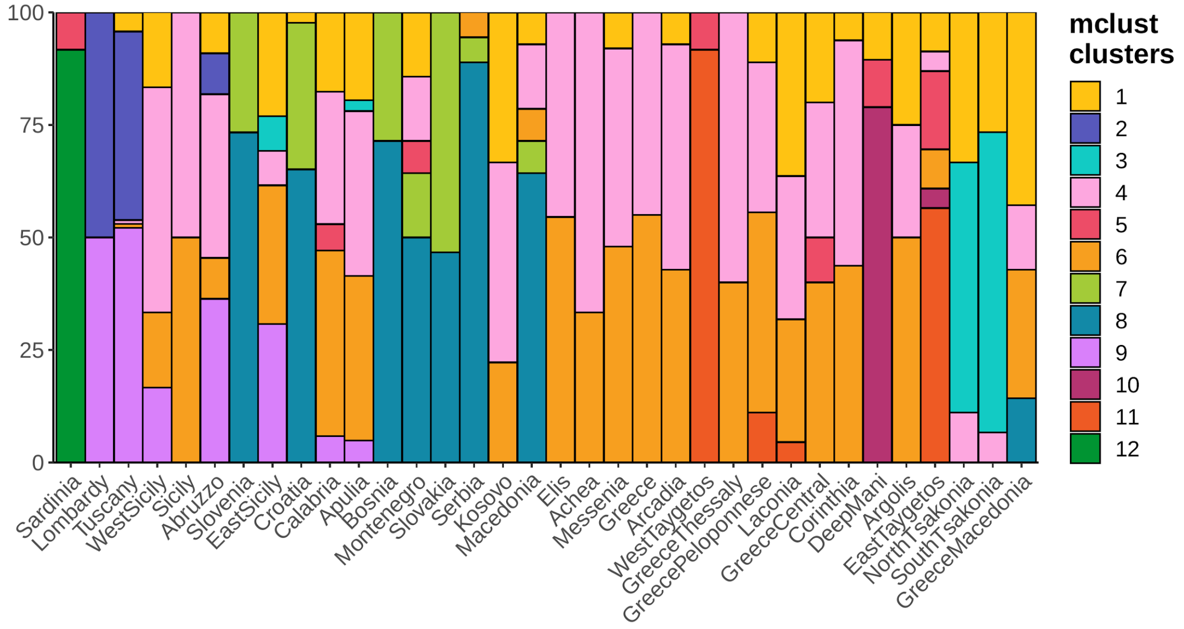


**Fig. S4**. **Genetic clusters.** The proportion of each bar represents the proportion of individuals who are grouped in each cluster by *mclust* analysis.

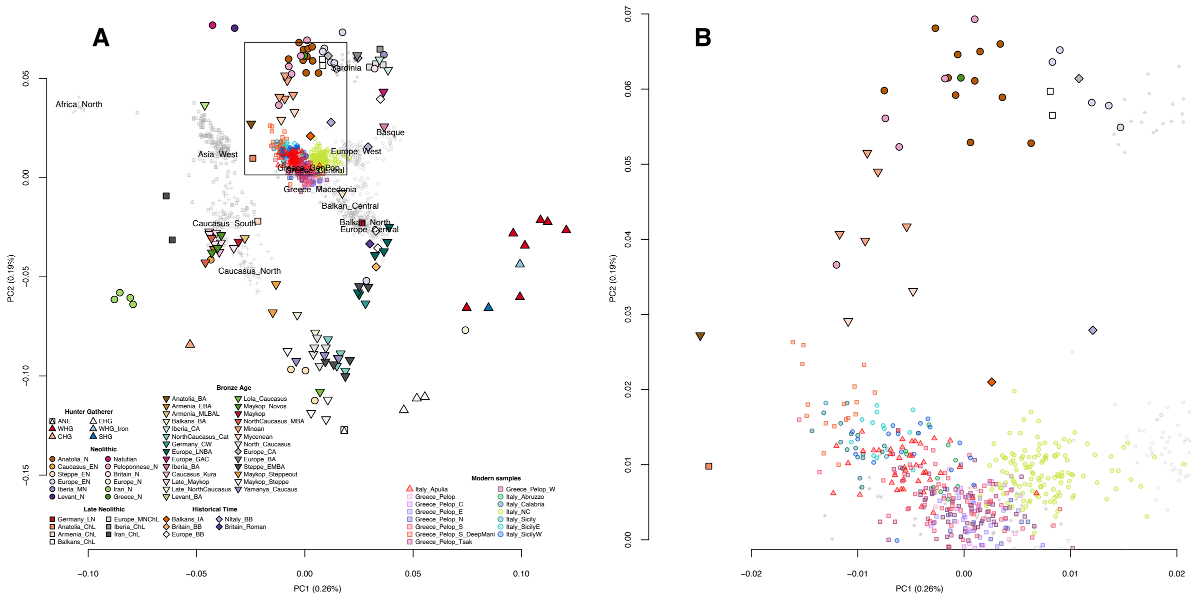

**Fig. S5**. **Ancient Eurasian genomes on European genetic variation.** A) PCA results from present-day Europeans (not black-bordered symbols) with ancient individuals projected on the first two components. Label of the population is shown for individuals who have been indicated with grey not-filled symbols and it is placed in correspondence of the median values of the first two components (PCs) of those individuals. B) Zoom of the PCA corresponding to the rectangle shown in panel A. Label codes can be found in Table S1.

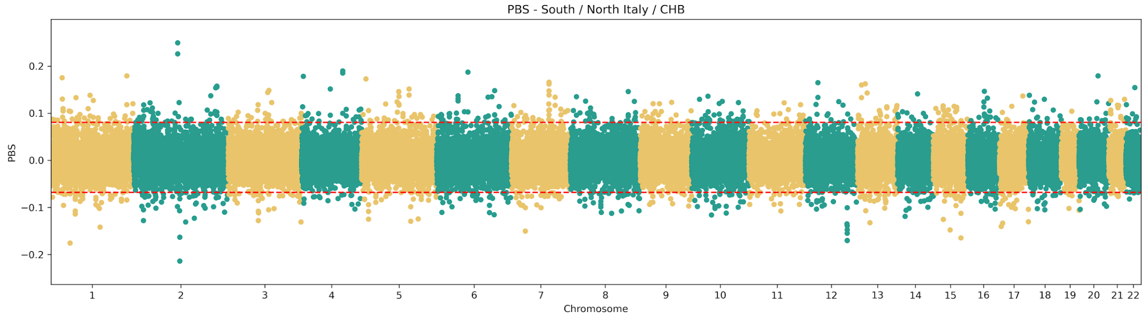


**Fig. S6**. **Manhattan plot for the PBS analysis between Southern and Northern Italy using CHB as outgroup.** All the loci analysed are reported as dots. Red dotted lines represent the top 0.05% of the statistic.

#### Supplementary Table (provided as Excel Tables)

**Table S1**. **Present-day and ancient samples used in this study.**

**Table S2**. **Highest *f4-outgroup* statistics for Southern Italian individuals with North and South Caucasus.** The test was performed in the form of f4(Mbuti, Caucasus, SE European, C European population). As a Caucasus populations, the Lezgins ethnic group (Behar et al., 2013, 2010) and present-day people inhabiting Georgia (Behar et al., 2013, 2010) were used.

**Table S3**. ***f4-outgroup* statistics for Steppe ancient source in OIG and present-day Northern Italian/Peloponnesian populations.** The test was performed in the form of f4(Mbuti, Steppe EMBA, OIG, Peloponnese/Lombardy).

**Table S4**. ***f4-outgroup* statistics to detect African sources in Southern Italy.** The test was performed in the form of f4(Mbuti, Yoruba, OIG, Lombardy).

**Table S5. Gene annotation of top 10 windows from PBS analysis. We extracted the top 10 positive and negative values of 5-SNP windows. Genetic coordinates (hg19) mean PBS value, the associated Z score and the genes encompassing the windows are reported.**
